## Supplementary Figure S1 for "Analysis of 8839 pan-primate retroviral LTR elements with regulatory functions during human embryogenesis reveals their global impacts on evolution of Modern Humans": List of abbreviations.pdf

HERVH, human endogenous retrovirus type H;

HERVL, human endogenous retrovirus type L;

LTR7, long terminal repeat 7;

MLT2A1, long terminal repeat MLT2A1;

MLT2A2, long terminal repeat MLT2A2;

GREAT, Genomic Regions Enrichment of Annotations Tool;

GSEA, gene set enrichment analyses;

NHP. Non-human primates;

MYA, million years;

GRD, genomic regulatory dominion;

GRN, genomic regulatory networks;

GRM, genomic regulatory module;

GRP, genomic regulatory pathways;

HSRS, human-specific regulatory sequences;

DEGs, differentially expressed genes;

ECA, extinct common ancestor;

lncRNA, long non-coding RNA;
